# Life history evolution of insects in response to climate variation: seasonal timing versus thermal physiology

**DOI:** 10.1101/2025.02.05.636251

**Authors:** Karl Gotthard, David Berger, Patrick Rohner

## Abstract

Climate adaptation in insects can proceed via responses in life history traits and their thermal plasticity, and through phenological shifts mediated by responses to photoperiodic cues (“photoperiodism”). While, experimental studies demonstrate evolutionary potential for both modes of adaptation, it remains unclear how evolution will unfold in natural populations, limiting our ability to predict how insects will respond to climate change. Here we review the literature and perform an analysis of published studies revealing that photoperiodism for diapause induction evolves predictably along latitude, with high-latitude populations entering diapause earlier. In contrast, although a few species showed clinal variation in life history and thermal plasticity, the direction of these clines were not consistent across taxa. This suggests that while insect life history and physiological adaptation to temperature can evolve, phenological shifts via evolution of photoperiodism is likely to be a more common and predictable response to future climate change.

## Introduction

Ecological and evolutionary processes influenced by climate variation are widely studied due to the urgent need to understand the impact of global climate change on biodiversity (3, 54, 114, 117, 148, 149, 153, 161). It is well established that global warming has already had strong ecological effects on biological communities by affecting the distribution, abundance, and phenology (seasonal timing) of many organisms, including insects (28, 41, 51, 53, 95, 115, 116, 148, 157). However, the mechanisms by which insects respond and adapt to climate change is less well understood (3, 23, 45, 58, 78). This presents a challenge not only for assessing future extinction risk of focal species, but also for predicting stability of species interactions and ecosystems as functions of alternative evolutionary strategies in response to a warming world (6, 24, 88). For instance, while changes in climate can lead to the evolution of physiological responses that mitigate environmental stress (45, 58), the evolution of phenology might offer an alternate route to adaptation (17, 20, 76) (Figure 1). While physiological and phenological responses may both facilitate adaptation, their effects on insect growth rates and distribution ranges are likely to differ. Here, we review the literature on insect life history evolution in response to variation in climate, with a particular focus on work that has explored genetic responses in both thermal plasticity and phenology via the use of photoperiodic cues (from hereon: “*photoperiodism*”).

**Figure 1:**
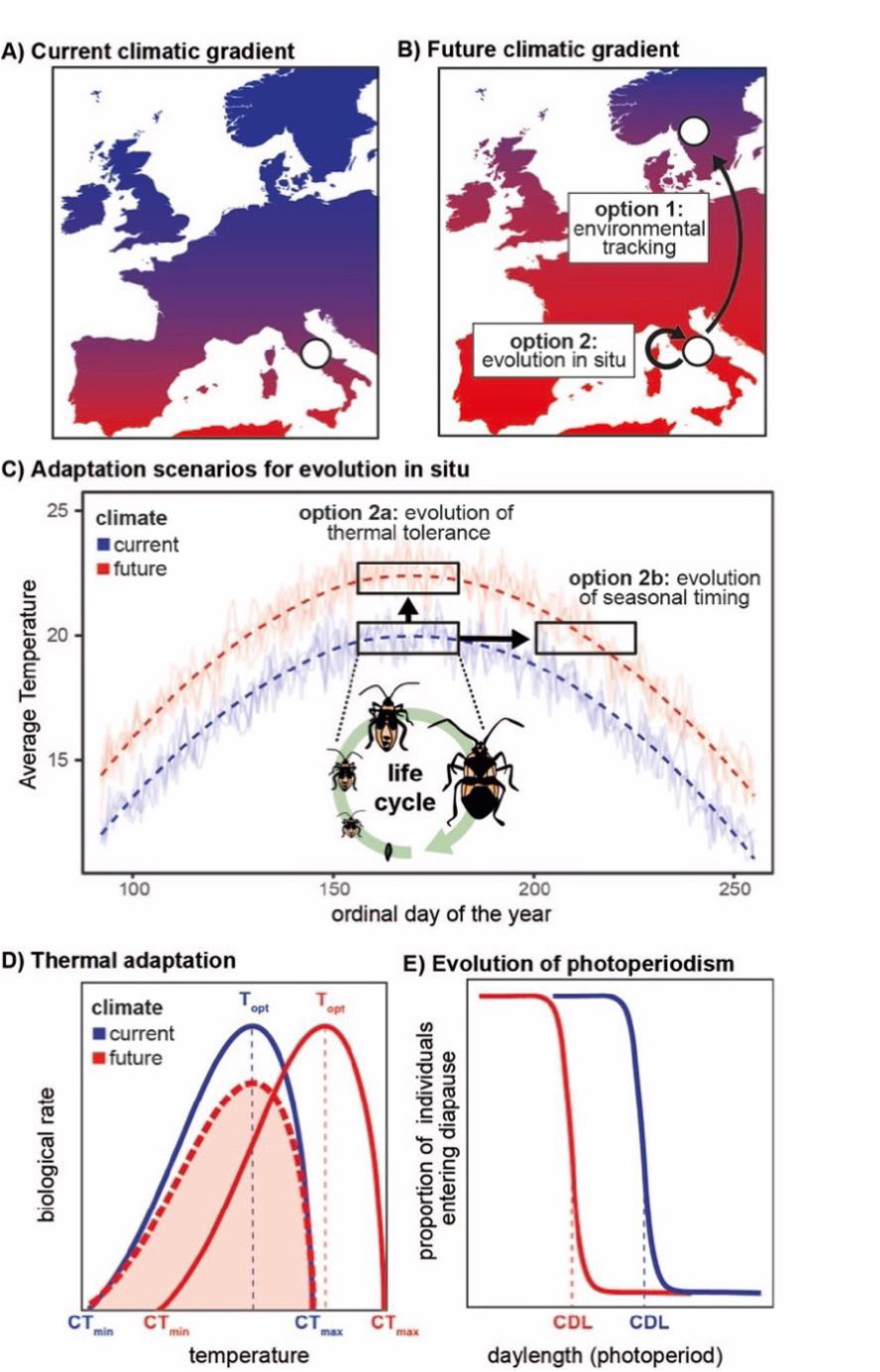
Insects have several mechanisms to respond to changing climates. A & B) Populations may track their preferred environment by shifting their geographic distribution range (option 1). Alternatively, they may remain in place and adapt to the new climatic conditions (option 2). C) This adaptation can occur through the evolution of thermal tolerance (option 2a) and/or changes in seasonal timing (option 2b). D) An organism’s thermal tolerance evolves through changes in the minimal (CT_min_), optimal (T_opt_), and/or maximal (CT_max_) temperature at which it can operate. As thermal tolerance is governed by trade-offs between performance at different temperatures, climate warming can result in evolutionary shifts in all these parameters (blue to red reaction norm). Note, however, that climate adaptation of growth and development also can occur through vertical shifts of the entire reaction norm (between blue and broken red reaction norm) so that growth is faster in northern populations that have less time and permissible thermal conditions to complete development, or so that growth slows down in future warming climates to counteract the direct effect of temperature on biological rates (so called “countergradient” variation). E) An organism can change its seasonal timing via the evolution of photoperiodism; by modification of how it uses and interprets daylength. Adaptation to climate warming by shifts or prolongations of the timing for growth and reproduction towards colder parts of the year (option 2b) is predicted to be associated with changes in the critical daylength (CDL) at which diapause is induced (blue to red reaction norm). Additionally, if species respond to warming climates by migrating closer to the poles (option 1), they also need to evolve changes in how they interpret daylength in order to time their life cycle. Importantly, the evolution of seasonal timing can relax selection for changes in thermal tolerance (and vice versa), and changes in seasonal timing may also require life history adjustments to accommodate changes in season length, such as overall increases or decreases in development rate and size at maturation, predicting correlated evolution of photoperiodism and life history.

### Physiological temperature adaptation in life-history traits

Life history theory sees the scheduling of life cycle events such as growth, maturation and reproduction as the result of strategic decisions made over an organism’s life (99, 111, 128, 140). Spatiotemporal variation in climate variables such as temperature, precipitation, and humidity are major sources of natural selection on life history strategies in general, and maybe in particular for ectotherms like insects (3, 23, 45, 58, 66, 111, 146). Virtually all physiological processes of insects are at least somewhat dependent on temperature, leading to abundant phenotypic plasticity in life history responses to ambient temperatures (3, 23, 111). For instance, decreased temperature often leads to plastic increases in adult body size in arthropods (the so-called temperature-size rule) (5, 59) and prolongs egg-to-adult development time (3, 23, 111). Nevertheless, theory predicts that an organism’s range of tolerated and preferred temperatures should evolve to match those experienced in its environment (3, 63), and indeed, there are often evolved species differences in the degree to which species are affected by temperature (3, 23, 111). Moreover, there is often noticeable genetic variation within species, indicating that thermal plasticity can evolve, sometimes even on ecological timescales (12, 52, 93). This suggests that natural selection may alter temperature dependence of important physiological and life history traits (3, 12, 38, 52, 97, 159), offering an avenue to adapt to climate change (Figure 1C-E).

### Seasonal adaptation in life-history via photoperiodism

Most organisms experience some type of seasonal variation in climate that also drive associated variation in biotic resources and interactions (22, 36, 146, 150). Seasonality is consequently a highly predictable form of temporal variation in climate and other ecological conditions (66, 146). Adaptation to seasonality requires that organisms can exploit the favorable seasons, avoid or mitigate the effects of the unfavorable seasons, and adequately time the transitions between respective stages in their life cycle (22). This is especially important for insects because most species use some kind of dormancy to withstand harsh conditions (36, 146, 150) (although there are notable examples of insects that migrate long distances to evade hazardous conditions (29, 146, 169)). In temperate insects, the typical adaptation to mitigate the effects of the unfavorable winter season is to enter diapause before the onset of cold conditions (i.e., a state of dormancy that arrests or greatly prolongs development). The individual then remains in diapause until there is no longer a risk of developing prematurely and finally restarts development and reproduction when the thermal conditions again become permissive (36, 83, 146, 160). Although variation in thermal conditions and biotic resources are likely to be the major sources of selection on life cycle timing, temperate insects have commonly evolved to use the photoperiod as a cue of seasonal change (36, 66, 146, 150). Photoperiod provides almost noise-free information on seasonal timing, allowing insects to appropriately schedule diapause. The sensitivity to photoperiod typically follows a threshold function where in certain photoperiods (typically long summer daylengths) individuals follow non-diapause (or ‘direct’) development, while in other photoperiods (typically shorter late summer/autumn daylengths) they instead enter diapause (76, 146). The inflection-point of this threshold function is referred to as the critical daylength (CDL) and is often used as a metric for assessing genetic variation in photoperiodism between populations or species (16, 22, 36, 76, 138, 146). This type of photoperiodic plasticity allows a given genotype to develop without a diapause in the warm summer months while still inducing diapause in individuals that are active later in the season. The facultative induction of diapause allows insect populations to produce multiple generations within a given year (multivoltinism) and genetic variation for CDL is often strongly associated with variation in voltinism (92, 138), and the length of the permissive season for growth and reproduction (76).

## Two modes of adaptation

Temperature variation during the favorable season is likely to exert strong selection on thermal plasticity of life history traits, whereas temperature variation at the end of the active season will exert selection on photoperiodic regulation of diapause induction. It is important to note that these two types of selection pressures interact with each other. On one hand, evolved responses in photoperiodism can shield the organism from unfavorable temperatures. On the other hand, evolved responses in thermal plasticity can in turn relax selection for shifts in phenology. Hence, we expect these two interacting modes of adaptation to exert correlational selection on the optimal life history strategy to mitigate climate variability (Figure 1C-E) (23, 45, 86, 168). However, the relative significance of selection on thermal physiology versus selection on photoperiodism is not well understood. Here we review how insect life histories have evolved in response to climate variation across both time and space by comparing cases where climate variation is expected to exert natural selection on life history traits and their thermal plasticity, as well as on photoperiodism for life cycle regulation.

### Evolution of thermal performance and photoperiodism in selection experiments

Experiments using artificial selection or experimental evolution to study the evolution of thermal reaction norms (i.e. thermal plasticity) have been performed in many species (3, 96). These experiments provide a proof-of-principle of how natural selection can shape thermal plasticity in life history. A recent meta-analysis of such experiments on ectotherms compared thermal reaction norms of evolved populations after thermal selection, to control (or ancestral) populations, allowing quantification of thermal adaptation (96). About a quarter of the studies included in the meta-analysis were done on insects. The results show a general positive response to thermal selection, where estimates of fitness in the assay temperature increased after selection. Moreover, thermal selection in the laboratory also had the capacity to change the shape of reaction norms so that adaptation to warmer temperatures generally led to lower fitness at colder temperatures (96). Although these effects were generally strongest in unicellular organisms, they still suggest that selection over relatively short timespans has capacity to alter the thermal sensitivity of life history traits and that there may be genetic trade-offs between performing at different thermal conditions.

Different types of artificial selection experiments that explore the evolutionary potential of photoperiodism for diapause induction and duration have been performed on species from all large insect orders (146). Typically, they show that these seasonal adaptations can change relatively rapidly in laboratory experiments (146). For instance, artificial selection on the photoperiodic induction of reproductive diapause in the fly *Drosophila montana* demonstrated a rapid change in critical daylength (77). Using a baseline population derived from a northern location in Finland, selection for eight generations where only non-diapausing individuals were allowed to breed under continuously shorter daylengths resulted in shorter CDL in all replicate lines compared to the control. These selected lines evolved a response to photoperiod that is typical for more southern populations of *D. montana*.

Similarly, bidirectional artificial selection for non-diapause and diapause development led to the evolution of photoperiodic plasticity for diapause induction in the North American mosquito *Wyeomyia smithii* after 12 generations (18), and in the chrysomelid beetle *Ophraella communa* after 5-8 generations (144).

In summary these examples demonstrate that both thermal reaction norms and photoperiodic reaction norms have the potential to evolve relatively rapidly in response to directional selection in the laboratory, suggesting that there is standing genetic variation present within natural populations that can fuel rapid evolutionary change in the wild.

### Life history evolution in response to recent climate change

Rising temperatures are altering patterns of seasonality and phenology — a trend expected to be further exacerbated in the foreseeable future (3). Against the backdrop of strong declines in insect abundances in recent decades (132, 157), there is increasing interest in the effects of climate on insect populations and their life histories (54, 161). However, to detect and estimate phenotypic evolution in natural populations it is necessary to measure heritable (i.e., ‘genetic’) changes in traits in the same populations repeatedly (at least twice) over several years or even decades.

In insects these types of studies are relatively rare as they require controlled common garden experiments to be replicated over extended time periods. Still, there are some notable examples, spearheaded by work on the North American Pitcher plant mosquito (*Wyeomyia smithii*) by Bradshaw and Holtzapfel (16). The authors demonstrated that the photoperiodic regulation of the life cycle, measured by the CDL for diapause induction in a range of latitudinally separated populations, showed a general genetic shift towards shorter CDLs, that are typical in more southern populations. This shift likely represents adaptive evolution in response to a warming climate as shorter CDLs result in a longer growth season and a later date for diapause induction. Surprisingly, this evolutionary change was detectable over only five years of evolution in the field (16). Similar results have since been found in other insects (44, 107, 108), indicating that changes in photoperiodism for life cycle regulation may constitute a common evolutionary response to climate change.

However, there have also been documented evolutionary changes in life history traits over time. For instance, the Yellow dung fly (*Scathophaga stercoraria*) shows genetically based reductions in body size (13), and a study on the Winter moth (*Operophtera brumata*) found a reduction in egg development rate (155). The latter example presumably represents a rare demonstration of how climate change has led to the evolution of thermal plasticity of development rate. Interestingly, this response also seems related to selection for a change in phenology (155) as several populations in the Netherlands have evolved a reduction in egg development rate in a range of relevant spring temperatures, most likely due to selection for maintaining synchronization of egg hatching with the budburst of its host plant, the Oak *(Quercus robur)*, in the now warmer spring. Perhaps the most extreme example comes from a recent study on fruit flies, showing evidence for rapid life history adaptation to climate fluctuations over a single season (spring to fall), corresponding to approximately 10 overlapping fly generations (129).

### Life history evolution in response to climate variation in space

Climate and seasonality vary greatly in space, and this variation is particularly predictable along latitudinal gradients (66). Studies of life history evolution in insects have a long tradition of exploring this type of latitudinal variation in climate (3, 14, 127, 137, 146). In many cases researchers have exploited this spatial variation to explore the likely effects of current climate change by the so-called “space-for-time” substitution (66).

There is a large body of literature that explores the thermal biology of life history evolution across insect species (3). In particular variation in thermal tolerance limits (i.e., the critical minimum and maximum temperatures, CT_min_ and CT_max_, respectively) across species and geographic locations are well-studied (2, 11, 57, 75, 142). Recent research has also highlighted the importance of studying corresponding measures of thermal tolerance in reproductive traits (40, 162), which may shed further light on adaptive potential and genetic constraints on climate adaptation in insect life history. A general pattern across insects (2, 57) and in ectotherms in general (11, 142) is that lower thermal limits show greater adaptive variation across climate regions at the species-level, compared to upper thermal limits. As a consequence, many researchers have converged on the conclusion that there may be limited scope for rapid evolution of CT_max_ in response to a warming world (11, 57, 75). Still, artificial selection experiments show that populations of the Fruit fly (*Drosophila melanogaster*) harbors additive genetic variation that allows successful laboratory evolution of increased heat tolerance, even if it may still be insufficient to keep up with the predicted increase in future temperatures (52). Moreover, as temperatures during the hottest parts of the year vary less across the typical geographic regions studied than do thermal minima, it is also possible that the relatively modest differentiation in CT_max_ can in part be explained by similar selection pressure across geographic regions (72). Studies exploring latitudinal variation in thermal tolerance within species provide a similar, albeit less strong, pattern as the between species comparisons (7, 9, 79, 80, 103, 125, 134, 136, 158, 167, 175). For example, within the limited data we reviewed for our quantitative analysis (see below), we found estimates of latitudinal variation in heat and cold tolerance for 4 and 8 dipterans, respectively. The observed clinal variation was not very consistent among species but perhaps slightly more so for cold tolerance, which tended to be higher in more northern populations. Interestingly, insects that have colonized urban environments show parallel evolution of both CT_max_ and CT_min_ (38, 97, 101), presumably due to local adaptation to the generally warmer urban environment. Together, this suggests that even though adaptive responses to natural selection on survival at thermal extremes do occur as insects colonize new climate zones, the rate of evolution may often be relatively slow.

There are also a large number of insect studies that have explored the evolution of development rates, body size, and their thermal plasticity across latitude (3, 14, 64, 73, 81, 98, 109, 112, 137, 139). However, predictions of how natural selection should affect development rate, size and their temperature dependencies are not always straightforward (15, 30, 59, 137). A common expectation is that because both mean temperatures and the length of insect growth seasons decrease with distance from the equator, natural selection should favor faster development rates (shorter development times) and thermal plasticity that maximizes growth in colder conditions in populations living at higher latitudes (81, 82, 127). This quite frequently observed pattern has been called “counter gradient variation” or “genetic compensation”, as genetic adaptation across latitude opposes the direct effect of temperature on plasticity in growth and development rates (32, 33, 48, 81, 82, 90). As a consequence of the shortened time for growth and development, it can be expected that adult size of insects should become smaller when moving away from the equator (127). However, physiological models assuming that rates of growth and development have different thermal plasticity predict that there are situations when insects should grow to larger sizes at colder latitudes (156, 163). Studies of latitudinal clines in body size within insect species have indeed found evidence for both these patterns (15, 81, 137). For instance, Shelomi (2012) reviewed a large set of both inter-and intraspecific latitudinal and altitudinal clines in insect body sizes, and while there were approximately equal numbers of positive and negative size clines, the most common pattern was the absence of a size cline altogether (137). Similar mixed patterns were found in a metanalysis of within-species size clines in arthropods that was mainly based on insects (59). Interestingly, this study found strong evidence that terrestrial taxa were more likely to show reductions in size with latitude, whereas aquatic taxa typically showed the opposite pattern. This result may be explained by the difference between water and air in how oxygen limitation affects both the plasticity and evolution of growth and development. Moreover, in the terrestrial taxa the pattern of voltinism influenced the latitudinal size clines as species with one or two generations per year decreased in size with latitude, while multivoltine species instead showed a weak positive size cline with latitude.

This suggests that as the length of the growth season increases towards the equator, species with strong latitudinal clines in voltinism (multivoltine) are under selection to produce additional generations, while in uni-and bivoltine species selection instead favors an increase in adult size via prolongation of development (59).

Because photoperiods change very predictably across latitudes, it is expected that natural selection should favor latitudinal variation in adaptations for seasonal timing in response to daylength. Indeed, there is a large number of examples demonstrating local adaptation in photoperiodism of insects, in particular in plasticity for diapause induction (35, 66, 76, 146). The most common pattern is that the critical daylength for diapause induction (CDL) increases with latitude across populations, which shows that more northern populations with a shorter growth season typically enter diapause earlier in year (76, 146). Latitudinal clines in CDL can evolve relatively rapidly in introduced species (10, 43, 146, 154), species colonizing urban areas (100), and in species expanding their ranges due to climate change (67).

The available evidence consequently suggests that life history traits such as development rates, body size and their thermal dependencies, as well as the photoperiodic plasticity for seasonal life cycle regulation do evolve across space and to some degree also with ongoing climate change. However, it is still unclear whether natural populations are more likely to adapt through changes in thermal plasticity, the adjustment of seasonal activity (through the evolution of photoperiodism), or both. Previous attempts to assess whether thermal-or photoperiodic adaptations are more likely to evolve in response to rapid climate change suggested that genetic changes in photoperiodism for seasonal life cycle regulation was a considerably more likely “first response” compared to thermal adaptation (17, 20). As these early reviews were based on the relatively limited set of then known examples and that selection due to climate change is likely to be multifaceted, other researchers have argued that this conclusion is premature (58). Unfortunately, well established examples of evolutionary change due to recent climate change are still rare, and for insects we have not found more than the handful of examples reviewed above. However, because of the large number of studies exploring evolution of photoperiodism, as well as life history traits and their thermal dependencies in response to climate variation across latitudes, it is possible to explore this question in the spatial dimension. Below, we first summarize and integrate the results of previously published studies to more formally quantify the amount of population differentiation in photoperiodism across latitude. We subsequently compare this to the corresponding latitudinal variation in life history traits and their thermal plasticity in the same set of species.

## Analysis of latitudinal clines in photoperiodism, life history, and thermal plasticity

Full details of the literature search and analyses can be found in the supplementary material. In brief, we looked for published common garden datasets that quantified latitudinal clines in critical daylength as well as means and thermal plasticity of the life history traits body size and development rate. The dataset on critical daylength was based on an initial data set generated by Joschinski and Bonte (2021) (76). We added additional data that has since been published and had at least 2 populations and 3 photoperiod treatments. Data on life history was collected based on an independent literature search that targeted species with photoperiodism data, again focusing on common garden studies that reported latitudinal clines. We calculated an index of thermal plasticity (i.e. how temperature affected a linear response in trait means) for those studies that included at least two temperature treatments to. Our final dataset included 308 populations from 24 insect species that had estimates of latitudinal clines in both critical daylength and development rate and/or body size. This included representatives of the orders Coleoptera (n = 5), Diptera (n = 5), Hemiptera (n = 3), Lepidoptera (n = 10), as well as one species of Neuroptera, from the original publications: (1, 4, 8, 14, 18, 27, 34, 42, 47, 55, 60–62, 67, 69, 70, 74, 85, 89, 92, 94, 104–106, 110, 113, 118, 119, 123, 126, 130, 131, 133, 135, 139, 143, 147, 151, 154, 164, 165, 171–174, 176).

We found that the vast majority of species showed an increase in CDL with latitude, indicating that populations closer to the poles have adapted to the generally longer summer photoperiods at these locations (Figures 2,3). Based on critical daylength estimates, we also calculated the mean ordinal day of the year at diapause onset at each population’s location. Average diapause onset showed a significant decrease with latitude, indicating that populations closer to the poles generally enter diapause earlier in the year. These findings are consistent with other studies (76) and indicate a strong and consistent evolutionary pattern of photoperiodism evolution across insect orders.

**Figure 2:**
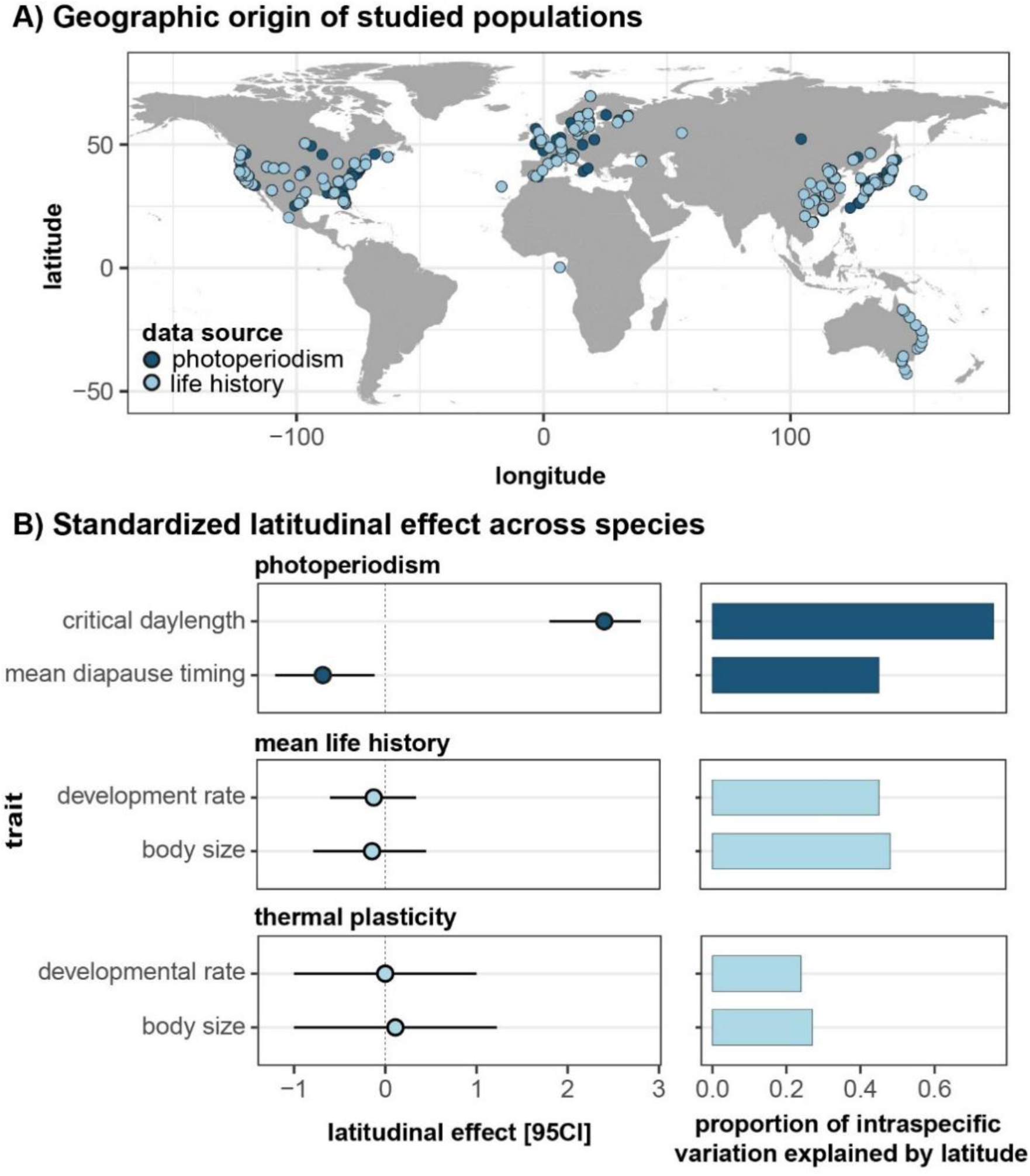
A) The geographical locations of all 308 populations from the 24 species that were included in the analysis of life history and photoperiodism. B) shows the global effects of latitude on photoperiodism, mean life history, and thermal plasticity in life history (means and corresponding 95% confidence limits). Only photoperiodism shows a strong and consistent clinal pattern across species. The standardized latitudinal effect refers to the global slope estimated in a Bayesian multilevel model divided by the standard deviation in the slope observed across species. Thus, this effect size estimate corresponds to a measure of repeatability across species. The horizontal bars to the right show the average proportion of population differentiation within each species that was explained by latitude (as measured by partial R^2^). Note that although life history traits did not show consistent clinal variation across species, latitude still explains a considerable fraction of the observed population-level variance in mean body size and development rate indicating evolution of latitudinal clines in opposing directions across species. Details are provided in the Supplementary material.

**Figure 3:**
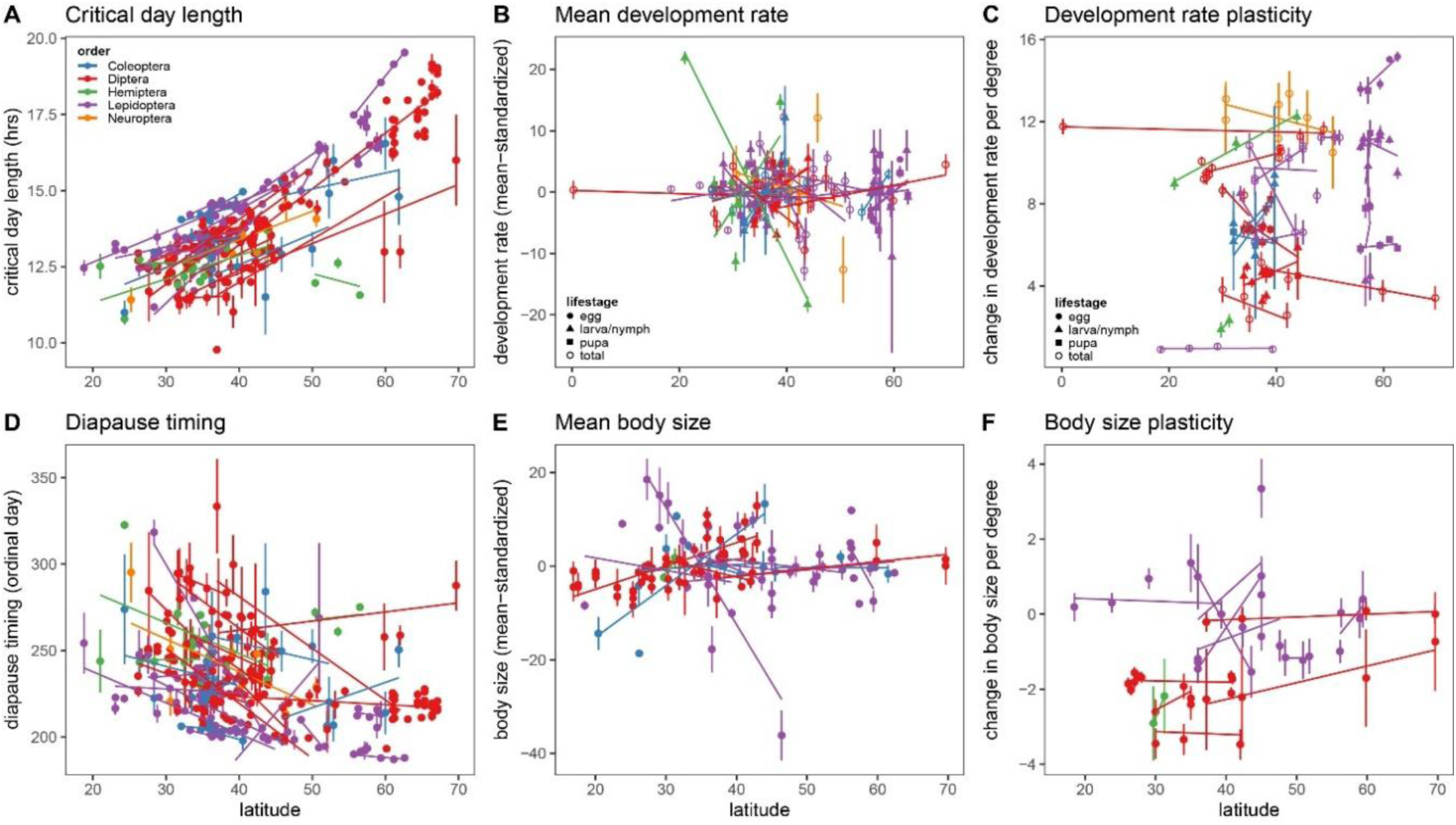
Population differentiation in photoperiodism, life history traits, and their thermal plasticity in relation to absolute latitude of sampled populations for each species (indicated by separate slopes). A) Critical day length increases consistently with latitude. In contrast, average life history (B, E) and life history plasticity (C, F) do not show consistent latitudinal patterns across species. D) The average day of the year at which the critical daylength for diapause is reached at each population’s location (“diapause timing”) decreases with latitude, indicating that diapause is induced at earlier dates at higher latitudes. The life stage in which development rates were estimated is specified in B and C. To facilitate comparison development rate and body size were standardized relative to the average trait value within each study and are expressed as percent change relative to this average. Thermal plasticity is expressed as the percent change in trait value (relative to the global average) per degree Celsius. Error bars indicate standard errors. Slopes are drawn separately per study in panel F; corresponding to 9 species with data from 12 studies.

In contrast, in the same set of species, we found no consistent latitudinal effect for any of the investigated life history traits (development rate, body size and the thermal plasticity of these traits). This finding contrasts with the notion that counter gradient adaptation in development rate is one of the main hypotheses for the evolution of latitudinal clines in life histories of insects. Importantly, however, there are many instances of life history clines in the data. In fact, latitude explained a considerable fraction of the observed population differentiation within species for mean development rate and body size (45% and 48%, respectively) (Figure 2B). For instance, the speckled wood butterfly (*Pararge aegeria*) and southwestern corn borer (*Diatraea grandiosella*) showed rather strong genetic decreases in development rate with latitude (47, 92, 143). However, patterns were inconsistent, and sometimes in opposing directions, across species (Figure 3) and we even found a couple of instances of inconsistent patterns within species. For example, in the Colorado potato beetle (*Leptinotarsa decemlineata*), development rate increases with latitude in European populations but decreased towards the pole in North America (69, 89). As latitudinal clines in body size often are considered to be results of a trade-off with development time (potentially also affected by shifts in voltinism), which showed no consistent latitudinal pattern here, the idiosyncratic responses in body size clines are perhaps less surprising. We also found no general pattern in thermal plasticity of body size or developmental rate. Although sample sizes were limited for those comparisons, most estimates of the latitudinal effect on thermal plasticity for individual species hoover around zero (Figures 2B and 3C, F). Taken together, these findings suggest that the mechanisms underpinning seasonal timing evolve predictably across orders. In contrast, while life history traits often do evolve, the patterns are not consistent across insects and remain hard to predict.

An important caveat is that there is still a relatively small number of species for which we were able to find both photoperiodism and life history data and we were not able to generate replicated measurements within species (except for a few species measured from more than one continent). This limits our ability to differentiate between evolved differences among species and spurious experimental effects, despite researchers’ best efforts to implement common garden rearings. However, by including only species where latitudinal variation in life history and CDL for diapause induction have been estimated, we were able to assess the relative likelihood of the two modes of adaptation in life history strategies. Although this reduced the number of species in our analysis, the results nevertheless revealed a very strong pattern in CDL and mean diapause onset and no consistent patterns in the other traits. In combination with the limited set of examples that have documented evolutionary changes due to recent climate change, it seems reasonable to assume that this higher predictability of the evolution of photoperiodism pertains also to the expected future changes in climate.

The difference in predictability of evolutionary change may have several explanations. Compared to life history traits, selection on CDL may simply be stronger and less dependent on other ecological circumstances that differ among species and geographic locations. For instance, variation in behavioral thermoregulation may affect the exposure to temperature variation, and moderate selection on thermal plasticity of life history traits (23). Along the same lines, and perhaps more pertinent to our analysis, it has been suggested that evolutionary changes in seasonal life cycle regulation may mitigate selection on thermal sensitivity (23) (Figure 1). This type of correlational selection would arise if selection typically favors a seasonal regulation that consistently exposes the active part of the life cycle to the time of the year that is most thermally similar at different locations, thereby reducing the difference in thermal selection across space; after all this is one of the main adaptive functions of insect diapause in general.

It is also possible that, compared to CDL, the amount of standing genetic variation is typically lower for thermal plasticity in life history traits, which would limit the response to selection. However, some studies indicate noticeable genetic variation for thermal plasticity of life history traits (26, 93), suggesting that it is unlikely to be the only explanation for the observed patterns. It is also possible that, even though thermal plasticity of life history traits harbor significant genetic variation, general differences in the genetic architecture of these different traits affect their evolvability. For instance, selection on life history traits and their thermal plasticity are likely to affect many genes with strong pleiotropic effects, which could lead to correlated evolution in other traits with maladaptive effects in the natural environment. For example, artificial selection causing adaptation to higher temperatures in a range of ectotherms led to correlated evolution of maladaptation at colder temperatures (96). As temperature has such a multifaceted influence on virtually all aspects of insect biology it is perhaps not surprising if a genetic change in one aspect of this temperature dependence also affects the temperature dependence of other important physiological processes. In a natural environment with highly variable thermal conditions such correlated effects are likely to lower the rate of evolution in response to a higher (or lower) mean temperature. In contrast, genes affecting photoperiodism may have less systemic effects and therefore be subject to less pleiotropic constraints. Indeed, several studies have found major effect loci that underpin intraspecific variation in diapause induction (46, 71, 84, 121, 122, 124, 145).

However, other studies suggest that photoperiodism is genetically highly complex (19, 21, 91, 124, 138) and some suggest that diapause induction and life history might be genetically correlated through linkage or pleiotropy (102).

Poleward range expansions are one of the most common effects of ongoing climate change (28, 115). Considering the general process that (re-)distributes insects across latitudes may provide yet another explanation for why genetic clines in CDL are more predictable than any of the other traits examined here. When insects colonize new locations across latitudes it is likely that their existing thermal adaptations are strong determinants of where they can establish successfully (68). This means that they may colonize new locations when thermal conditions have changed enough to allow positive population growth. If so, selection for rapid thermal adaptation may initially be comparatively weak in these newly colonized locations. In contrast, all expansions across latitudes will inevitably mean that they are exposed to a novel regime of photoperiodic variation and if a non-optimal CDL does not preclude establishment, the novel environment will consistently exert selection on photoperiodic plasticity and CDL (67). The latitudinal snapshots represented in our collated data may be a reflection of this process.

## Conclusions

Several studies indicate that insects can relatively rapidly evolve in response to climate change, either through the evolution of thermal physiology or shifts in seasonal activity. Our review suggests that changes in photoperiodism, that is, the timing of life history transitions via use of photoperiod cues, is a predictable way in which various insect species adapt to climate variability. In contrast, while changes in life history traits often evolve, there is little support for a general pattern that holds broadly across species. In the context of climate adaptation, evolution of seasonal timing consequently emerges as a more predictable factor than physiological adaptations to thermal conditions. As such, our updated review and more formal analysis of life history evolution in response to climate variation gives credence to the statement of Bradshaw and Holzapfel (2008): it’s seasonal timing that matters.

Because the evolution of photoperiodism (or lack thereof) can lead to phenological mismatches and affect species interactions and ecosystem functioning, these evolutionary changes may have major impacts on the structure and resilience of natural systems. Our review thus offers some hope regarding our ability to predict shifts in insect phenology and its associated consequences in future climates. However, it should be noted that obligate univoltine as well as tropical insects are less likely to rely on photoperiodic plasticity for seasonal life cycle regulation and diapause induction. These insects may instead be more dependent on thermal variation also for life cycle regulation and potentially be more likely to show predictable clines in life history and thermal plasticity in relation to spatial variation in climate (49, 50). Moreover, despite relatively rapid evolution of seasonal life cycle regulation, many insect populations will ultimately face significantly hotter conditions during important parts of their life cycles. Notable in this regard, our analysis is heavily biased towards terrestrial species with good capacity to disperse, whereas aquatic taxa and those with generally more stationary lifestyles are not represented. Such taxa may be particularly vulnerable to climate change (56). The comparatively slow evolution and limited scope for adaptive acclimation of upper thermal limits shown in many insects is therefore clearly worrying (11, 57, 75, 120, 166). It should also be noted that the analyzed data suffers from the general sampling bias in studies of biodiversity, with a strong underrepresentation of not only the tropics but also the southern hemisphere (25, 65) (Fig. 2A). Because the distribution of biodiversity as well as current and future climate niches are geographically heterogenous, future forecasting will need to account for the fact that climate change itself is expected to progress at different pace across the globe (31, 39, 170).

Predicting how climate change will impact insect life history is crucial to mitigating both biodiversity loss and pest outbreaks (37, 53, 88). Current statistical projection models indicate that climate warming will lead to deterministic changes in insect life history via temperature’s effects on biological rates (39) leading to range expansions and increases in abundance in temperate regions, but population decline in the tropics (37). However, these projections may have poor accuracy as they do not factor in evolutionary responses. For example, a recent study on the beetle pest *Callosobruchus maculatus* showed that less than a decade of life history adaptation at hot temperature in the laboratory almost doubled its predicted costs to agriculture compared to simple statistical projections (24). The unpredictability of observed life history evolution in the species analyzed here adds to the notion that forecasting responses in insect growth rates and distribution ranges will be a significant challenge (87). Our review highlights four critical areas for future work that may help tackle this problem. First, the number of species in which clines in diapause induction, life history, and various physiological responses have been tested in a common garden experiment remains small, taxonomically biased, and rarely replicated within species. Future research should aim at closing this gap. One possibility here is also to study variation along altitude, which typically replicates climate variation at a smaller spatial scale compared to latitudes. For instance, a recent study of altitudinal variation among populations of burying beetles found that while photoperiodism showed systematic population differentiation, there was no similar effect on thermal plasticity in life history (152), much in agreement with the general pattern in our multispecies analysis across latitudes. Second, there is a need for more studies that investigate how natural populations adapt in situ to changing climates (13, 16, 44, 107, 108, 129, 155). Although these longitudinal studies suffer from a variety of experimental challenges, they provide important insights into climate-induced life history evolution in the wild (141). Third, studying the genetic architecture of photoperiodism, life history, and thermal physiology will provide critical insights into the evolvability of natural populations - at the moment it remains unclear if the number of loci and their pleiotropic relationships differ qualitatively between thermal plasticity in life history and seasonal timing traits, and how important genetic constraints are in explaining the observed patterns. Finally, studies that characterize genetic (co)variation in photoperiodism and life history traits within and between natural populations should ideally combine these estimates with measures of their fitness consequences, as this will offer insights into the role of correlational selection in driving the observed clinal patterns. It is clear that these suggested research strategies will require major effort and skill in combination with significant funding, but given the magnitude of expected effects of climate change for both natural environments and human well-being, it is likely to be a valuable investment.

## Acknowledgements

This research was supported by funding from the Swedish research council (VR 2021-04258 to KG), the Swedish research council FORMAS (2022-01117 to DB). We would like to thank David Holway for helpful comments on earlier versions of this manuscript.

