## Supplementary Information for "Life history evolution of insects in response to climate variation: seasonal timing versus thermal physiology"

### Methods

*Literature search*

We extracted data on latitudinal clines in the critical day length for diapause induction (from hereon: "photoperiodism") from insect species from Joschinski and Bonte (21). In total, 433 reaction norms of 49 insect species had been scored for photoperiodism. In addition, we complemented this dataset with our own search of more recent studies, published from 2019 and onwards, applying the following search terms to the Web of Science Core Collection:

TS = *(altitude OR latitude OR latitudinal OR  cline)) AND (common garden OR adaptation OR adaptive OR evolution OR genetic)) AND (temperature OR photoperiod OR phenology OR seasonality)) AND (diapause OR life history OR development OR growth)) AND (insect OR Holometabola OR Endopterygota OR Orthoptera OR Diptera OR Odonata OR Hemiptera OR Hymenoptera OR Coleoptera OR Lepidoptera OR Trichoptera OR Mecoptera OR Siphonaptera OR Neuroptera OR Megaloptera OR Raphidioptera OR Strepsiptera OR Ephemeroptera OR Dermaptera OR Plecoptera OR Phasmatodea OR Mantodea OR Blattodea OR Thysanoptera OR Isoptera)*.

Including the major insect orders increased the total number of papers. To assess latitudinal variation in photoperiodism, we only considered common garden studies that included at least two populations and at least three photoperiods. This resulted in another 10 studies, with the total dataset on photoperiodism consisting of 67 studies on 58 species and a total of 510 geographically separated populations.

All species for which there were latitudinal clines in photoperiodism were then scanned for studies on latitudinal clines in life-history and thermal plasticity. We only included studies that measured at least two populations from different latitudes using a common garden design, so that the compared populations were measured in the same temperature and photoperiod. For species where several studies existed measuring the same traits, we picked the study that measured populations with the closest geographical distance to those measured for photoperiodism, unless the number of populations and temperature treatments were low (<3 populations and <2 temperatures) in which case we included more than one study if available.

After the full scan, we found 58 studies that fulfilled our criteria. However, only those studies measuring development rate (n = 30) or body size (n = 21), were judged to be numerous enough to be considered for analysis. We aimed at including studies which measured traits at more than one temperature so that we could calculate a measure of thermal plasticity (i.e. a response in population mean to temperature), although only 17 (development rate) and 9 (body size) studies included more than one temperature. We were not able to find a sufficient number of studies that investigated full thermal performance curves to consider them for our analyses. Our final dataset included 24 species that had overlapping estimates for latitudinal clines in critical daylength (CDL) and life history (see table S1).

*Estimating critical photoperiod and mean diapause timing in ordinal days*

Following Joschinski and Bonte (21), we extracted the proportion of individuals entering diapause at a given photoperiod and latitude from all studies, along with sample size. In cases where exact treatment-specific sample sizes were not reported, we divided the total number of individuals reported by the total number of treatments (thus assuming an even sample size for all treatment groups).

While changes in day length at a single location is a good indication of photoperiodism, it is not a good general indicator of phenology relative to other populations because daylengths depend on latitude. A change in CDL across populations thus does not necessarily result in a difference in the ordinal day of the year when days are long (or short) enough to initiate diapause. To generate a better measure of phenology across the globe, we calculated the ordinal day of the year at which a population's CDL would initiate diapause at the location where it was originally collected (using the R-package *geosphere* (13)). Some of the photoperiod treatments applied in the various studies were outside the range of naturally occurring photoperiods at the location where the population was originally collected. Following Joschinski and Bonte (21), we set these unnaturally long or short photoperiods to the daylength at that latitude at midsummer or midwinter, respectively.

To estimate critical daylength (CDL) and the ordinal day of the year where diapause is induced (mean diapause timing), we fitted a logistic model as:

$$f_x = \frac{d-c}{1+\exp\left(b(x-e)\right)} + c \qquad \text{Eq. 1}$$

Where *c* and *d* are the lower and upper asymptotes, *b* is slope at the inflection point, and *e* is the inflection point (i.e., the critical day length or mean diapause timing). This model was applied to each population with available diapause induction data, replicating the approach by Joschinski and Bonte (21). In brief, we ran Markov chain simulations for reaction norms using the R-package *rjags* (33). When available, observations were weighed by the sample size within each sample. In those cases where sample sizes were not available, we assumed equal weights. Four independent chains were run for 11,000 iterations each. The first 1,000 iterations of each chain were discarded as burn-in. We used uninformative priors for all four parameters; see Joschinski and Bonte (21). Parameter estimation for *e* was constrained within the range of photoperiods (or ordinal days) available for each population. Standard errors were calculated based on the posterior distribution. In some populations of *Zygaena trifolii* and *Scathophaga stercoraria*, this approach led to biologically unreliable estimates of critical day lengths because either all or very few individuals entered diapause. For these populations, we set the longest or shortest photoperiod included in the study as the critical photoperiod value and used the average standard errors of the other populations as an estimate of the standard error, noting that this approach produced conservative estimates of the strength of the estimated clines.

*Effect size calculations for life-history traits*

Body size estimates that were measured on a linear scale or as an area were transformed by cubic and 3/2 power-transformation prior to analysis to be comparable to those studies reporting mass. We added a moderator variable to analyses on body size stating whether size was reported as mass or by a linear measure. Development time was transformed into a rate (1/x) to linearize its relationship with temperature. Some studies recorded development rates of several life stages separately, and we therefore added a moderator variable including this information (egg, larval, pupal, or total development).

Within each study and combination of assay temperature and trait combination, we standardized reported population means and standard errors by dividing the estimates with the mean across all study populations measured at the same temperature, and multiplied by 100, thus giving the population- and temperature-specific deviation in trait mean in percent relative to the global average for a given temperature. This approach ensures that differences between populations are placed on relative scale so that general effects of temperature on trait means (e.g. much higher development rates at warm temperatures) do not a priori contribute to larger or smaller population differences. We then calculated the trait mean, $\bar{x}$, for population *j*, as:

$$\overline{x_j} = \frac{\sum_i^n x_{j,i}}{n} \quad \text{Eq. 2a}$$

where $x_{j,i}$ is the value for population $j$, at temperature $i$, which is then averaged over the total number of experimental temperatures, $n$. We approximated the standard error of this mean as:

$$SE_{\overline{x_j}} = \frac{\sum_i^n SE_{x_{j,i}}/n}{\sqrt{n}} \quad \text{Eq. 2b}$$

Our estimate of a population's thermal plasticity for trait $x$ was calculated as the slope of change in trait per degree Celsius:

$$P_{j,x} = \frac{x_{j,t_{max}} - x_{j,t_{min}}}{t_{max} - t_{min}} * \frac{100}{\bar{\bar{x}}} \quad \text{Eq. 3a}$$

Where $t_{max}$ and $t_{min}$ is warmest and coldest temperature used in the study. Here, $x_{j,t_{max}}$ and $x_{j,t_{min}}$ were standardized by dividing by the grand mean, $\bar{\bar{x}}$, taken over all populations and temperatures $t_{min}$ to $t_{max}$, and then multiplied by 100. The thermal plasticity is thus given as a slope reporting the percent change in trait mean of population $j$ relative to the overall mean, per degree Celsius. The standard error for the thermal sensitivity was calculated as:

$$SE_{P_{x,j}} = \frac{\sqrt{SE^2_{x,j_{min}} + SE^2_{x,j_{max}}}}{t_{max} - t_{min}} * \frac{100}{\bar{\bar{x}}} \quad \text{Eq. 3b}$$

For one study, the coldest temperature caused some species to not survive. Similarly, for two studies, the hottest temperature caused all populations to show strong non-linear declines in trait values. In both these cases we used the less stressful temperature for the calculations (i.e., $t_{min+1}$ and $t_{max-1}$, respectively).

For three studies (four estimates), we could not retrieve standard errors. For these cases we assumed that the standard errors were equal to the mean error calculated over the other studies in the dataset including the same trait and number of temperature treatments.

*Analyses*

For all analyses, we chose to classify entries of species occurring on separate continents as separate "species", given that these entries likely represent independent instances of (potential) latitudinal adaptation. This occurred for three cases: the fruit fly *Drosophila melanogaster* (Europe and Australia), the Colorado potato beetle *Leptinotarsa decemlineata* (North America and Europe), and the cabbage butterfly *Pieris rapae* (North America and Asia).

We estimated effects in Bayesian mixed effect models using the *brms* package (Bürkner 2017) for R. We first estimated a global effect of latitude on critical daylength, mean diapause timing, population mean development rate and body size, as well as thermal plasticity in development rate and size, accounting for variance in the effect of latitude between species.

$$y_{\bar{x}} = latitude + (latitude|species) + Error_{pop} + Error_{within} \qquad \text{Eq. 4a}$$

where $Error_{pop}$ and $Error_{within}$ designate residual variation between and within populations, respectively. We similarly estimated the global effect of latitude on thermal plasticity in body size and development rate, with the addition of adding an identifier variable (ID) accounting for variation in plasticity stemming from study/species differences, and the type of estimate for development rate (egg, larval, pupal or total) and body size (mass or linear).

$$y_{P_x} = ID + \ latitude + (latitude|species) + Error_{pop} + Error_{within} \qquad \text{Eq. 4b}$$

Note that adding the ID variable was not necessary for the analysis of average development rate and body size (above, Eq. 4a), as this variation was accounted for and removed by the mean-standardization of trait values. Note also that our analyses cannot readily distinguish between variation in trait means stemming from true species differences and those stemming from experimental methodology (e.g. choice of thermal range or food source) as most species were represented by a single study.

We also estimated species-specific latitudinal slopes by modelling these as fixed effects. We again note the lack of replication on the species level, but we find it less likely that the latitudinal effect within each species would be strongly affected by methodological issues.

$$y_{\bar{x}} = latitude{:}species + Error_{pop} + Error_{within} \qquad \text{Eq. 5a}$$

$$y_{P_x} = ID + \ latitude{:}species + Error_{pop} + Error_{within} \qquad \text{Eq. 5b}$$

For these latter two models, we quantified the partial $R^2$-value for how much of the intraspecific (between-population) variation within each species that was accounted for by latitude, following Gelman, et al. (9).

All models were run in brms using default priors for fixed and random effects, setting sigma = TRUE. To compare effect sizes across traits, we divided the grand mean latitudinal slope for each trait (Eqs. 4a,b) by the standard deviation in the slope-estimates across all species. Note that this effect size thus corresponds to a form of repeatability, reporting how large the observed effect is relative to the variation among species.

### *Results*

We find no evidence for global effects of latitude on mean development rate (b: -0.13 [--0.61, 0.34] 95% CI) or body size (0.11 [-1.00, 1.22]). Similarly, there was no global effect of latitude on thermal plasticity of development rate (0.00 [-0.08, 0.70]), nor body size (0.11[-1.00, 1.22]). On the contrary, there was a very strong and significant global effect of latitude on critical day length (2.4 [1.80, 2.80]). We also found a significant decrease in mean diapause timing with latitude (-0.68 [-1.21, -0.12]). Species-specific clines explained a large fraction of the intraspecific variation in critical day length ($R^2$ = 0.76) but less so for mean diapause timing ($R^2$ = 0.45).

Even in the absence of general latitudinal patterns in life history, species-specific adaptation along latitude can still be prominent. In our models estimating species specific latitudinal slopes, there were several cases showing significant clines for all traits along latitude. This was especially true for clines in mean trait values (Supplementary Fig. 1a), but less so for clines in thermal plasticity (Supplementary Fig. 1b). Indeed, latitude explained more of the intraspecific variation in population means ($R^2$ for development rate = 0.45; body size = 0.48) compared to intraspecific variation in thermal plasticity ($R^2$ for development rate = 0.21; body size = 0.27). This suggests that latitude has a strong effect on intraspecific variation in life history, but that these effects are species-specific and do not hold generality across species.

### Figures and tables

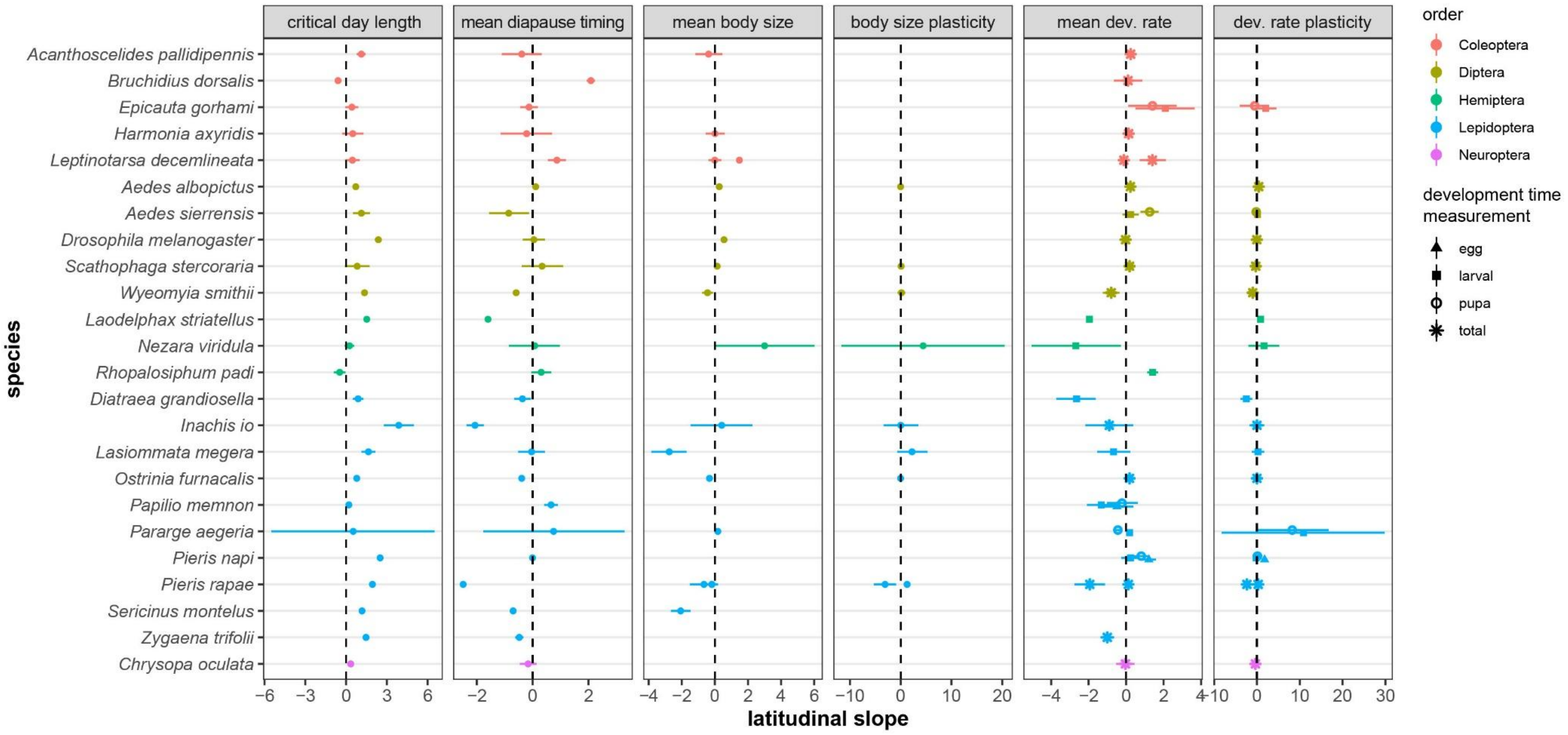


**Supplementary figure 1:** Species-specific slope coefficients (Means and 95%CIs based on Bayesian posterior estimates) for latitudinal patterns across populations examined under common garden conditions (estimated from models 5A and 5B). Positive values indicate that trait values increase at higher latitudes. To compare effect sizes across traits, the grand mean latitudinal slope for each trait (Eqs. 4a,b) was divided by the standard deviation in the slope observed across the species-specific means. In contrast to critical daylength, there was no consistent effect of latitude on any of the other traits when estimated across species.

**Table S1:** List of sources used to extract data for latitudinal effects on photoperiodism, life history, and thermal plasticity in life history

| Species | Order | Source of photoperiodism data | Source of life history and/or thermal plasticity data |
|---|---|---|---|
| *Acanthoscelides pallidipennis* | Coleoptera | (37) | (38) |
| *Aedes albopictus* | Diptera | (26, 45) | (2) |
| *Aedes sierrensis* | Diptera | (20) | (8) |
| *Bruchidius dorsalis* | Coleoptera | (22) | (22) |
| *Chrysopa oculata* | Neuroptera | (28) | (28) |
| *Diatraea grandiosella* | Lepidoptera | (42) | (42) |
| *Drosophila melanogaster* | Diptera | (31) | (19, 44) |
| *Epicauta gorhami* | Coleoptera | (43) | (43) |
| *Harmonia axyridis* | Coleoptera | (36) | (46) |
| *Inachis io* | Lepidoptera | (34) | (34) |
| *Laodelphax striatellus* | Hemiptera | (29) | (14) |
| *Lasiommata megera* | Lepidoptera | (17) | (17) |
| *Leptinotarsa decemlineata* | Coleoptera | (23) | (15, 18, 23) |
| *Nezara viridula* | Hemiptera | (27) | (7) |
| *Ostrinia furnacalis* | Lepidoptera | (16) | (49) |
| *Papilio memnon* | Lepidoptera | (50) | (51) |
| *Pararge aegeria* | Lepidoptera | (24) | (1, 11, 24) |
| *Pieris napi* | Lepidoptera | (41) | (41) |
| *Pieris rapae* | Lepidoptera | (12) | (10, 30, 40) |
| *Rhopalosiphum padi* | Hemiptera | (25) | (32) |
| *Scathophaga stercoraria* | Diptera | (5, 39) | (3, 5) |
| *Sericinus montelus* | Lepidoptera | (47) | (52) |
| *Wyeomyia smithii* | Diptera | (6) | (35) |
| *Zygaena trifolii* | Lepidoptera | (48) | (48) |

**Table S2:** Species-specific slope coefficients (means and standard errors based on Bayesian posterior estimates) for latitudinal patterns across populations examined under common garden conditions (estimated from models 5A and 5B). Positive values indicate that trait values increase at higher latitudes. To compare effect sizes across traits, the grand mean latitudinal slope for each trait (Eqs. 4a,b) was divided by the standard deviation in the slope observed across the species-specific means.

| Trait | Order | Genus | Species | Region | Trait modifier | Latitudinal slope | SE |
|---|---|---|---|---|---|---|---|
| Critical daylength | Coleoptera | *Acanthoscelides* | *pallidipennis* | Asia | | 1.123 | 0.168 |
| Critical daylength | Coleoptera | *Bruchidius* | *dorsalis* | Asia | | -0.592 | 0.071 |
| Critical daylength | Coleoptera | *Epicauta* | *gorhami* | Asia | | 0.426 | 0.247 |
| Critical daylength | Coleoptera | *Harmonia* | *axyridis* | Europe | | 0.491 | 0.401 |
| Critical daylength | Coleoptera | *Leptinotarsa* | *decemlineata* | Europe | | 0.465 | 0.278 |
| Critical daylength | Diptera | *Aedes* | *albopictus* | NorthAmerica | | 0.714 | 0.006 |
| Critical daylength | Diptera | *Aedes* | *sierrensis* | NorthAmerica | | 1.126 | 0.314 |
| Critical daylength | Diptera | *Drosophila* | *melanogaster* | Europe | | 2.373 | 0.13 |
| Critical daylength | Diptera | *Scathophaga* | *stercoraria* | Europe | | 0.819 | 0.465 |
| Critical daylength | Diptera | *Wyeomyia* | *smithii* | NorthAmerica | | 1.357 | 0.008 |
| Critical daylength | Hemiptera | *Laodelphax* | *striatellus* | Asia | | 1.522 | 0.025 |
| Critical daylength | Hemiptera | *Nezara* | *viridula* | Asia | | 0.255 | 0.179 |
| Critical daylength | Hemiptera | *Rhopalosiphum* | *padi* | Europe | | -0.472 | 0.218 |
| Critical daylength | Lepidoptera | *Diatraea* | *grandiosella* | NorthAmerica | | 0.885 | 0.199 |
| Critical daylength | Lepidoptera | *Inachis* | *io* | Europe | | 3.873 | 0.567 |
| Critical daylength | Lepidoptera | *Lasiommata* | *megera* | Europe | | 1.647 | 0.263 |
| Critical daylength | Lepidoptera | *Ostrinia* | *furnacalis* | Asia | | 0.786 | 0.038 |
| Critical daylength | Lepidoptera | *Papilio* | *memnon* | Asia | | 0.205 | 0.076 |
| Critical daylength | Lepidoptera | *Pararge* | *aegeria* | Europe | | 0.522 | 3.078 |
| Critical daylength | Lepidoptera | *Pieris* | *napi* | Europe | | 2.505 | 0.102 |
| Critical daylength | Lepidoptera | *Pieris* | *rapae* | Asia | | 1.939 | 0.031 |
| Critical daylength | Lepidoptera | *Sericinus* | *montelus* | Asia | | 1.172 | 0.032 |

| | | | | | | | |
|---|---|---|---|---|---|---|---|
| Critical daylength | Lepidoptera | *Zygaena* | *trifolii* | Europe | | 1.465 | 0.087 |
| Critical daylength | Neuroptera | *Chrysopa* | *oculata* | NorthAmerica | | 0.343 | 0.098 |
| Mean diapause timing | Coleoptera | *Acanthoscelides* | *pallidipennis* | Asia | | -0.391 | 0.367 |
| Mean diapause timing | Coleoptera | *Bruchidius* | *dorsalis* | Asia | | 2.087 | 0.076 |
| Mean diapause timing | Coleoptera | *Epicauta* | *gorhami* | Asia | | -0.13 | 0.163 |
| Mean diapause timing | Coleoptera | *Harmonia* | *axyridis* | Europe | | -0.216 | 0.473 |
| Mean diapause timing | Coleoptera | *Leptinotarsa* | *decemlineata* | Europe | | 0.869 | 0.165 |
| Mean diapause timing | Diptera | *Aedes* | *albopictus* | NorthAmerica | | 0.107 | 0.009 |
| Mean diapause timing | Diptera | *Aedes* | *sierrensis* | NorthAmerica | | -0.858 | 0.367 |
| Mean diapause timing | Diptera | *Drosophila* | *melanogaster* | Europe | | 0.044 | 0.201 |
| Mean diapause timing | Diptera | *Scathophaga* | *stercoraria* | Europe | | 0.337 | 0.379 |
| Mean diapause timing | Diptera | *Wyeomyia* | *smithii* | NorthAmerica | | -0.592 | 0.006 |
| Mean diapause timing | Hemiptera | *Laodelphax* | *striatellus* | Asia | | -1.6 | 0.026 |
| Mean diapause timing | Hemiptera | *Nezara* | *viridula* | Asia | | 0.072 | 0.469 |
| Mean diapause timing | Hemiptera | *Rhopalosiphum* | *padi* | Europe | | 0.307 | 0.184 |
| Mean diapause timing | Lepidoptera | *Diatraea* | *grandiosella* | NorthAmerica | | -0.367 | 0.153 |
| Mean diapause timing | Lepidoptera | *Inachis* | *io* | Europe | | -2.065 | 0.162 |
| Mean diapause timing | Lepidoptera | *Lasiommata* | *megera* | Europe | | -0.038 | 0.247 |
| Mean diapause timing | Lepidoptera | *Ostrinia* | *furnacalis* | Asia | | -0.393 | 0.023 |
| Mean diapause timing | Lepidoptera | *Papilio* | *memnon* | Asia | | 0.66 | 0.125 |
| Mean diapause timing | Lepidoptera | *Pararge* | *aegeria* | Europe | | 0.751 | 1.278 |
| Mean diapause timing | Lepidoptera | *Pieris* | *napi* | Europe | | -0.005 | 0.023 |
| Mean diapause timing | Lepidoptera | *Pieris* | *rapae* | Asia | | -2.491 | 0.033 |
| Mean diapause timing | Lepidoptera | *Sericinus* | *montelus* | Asia | | -0.7 | 0.046 |
| Mean diapause timing | Lepidoptera | *Zygaena* | *trifolii* | Europe | | -0.48 | 0.076 |
| Mean diapause timing | Neuroptera | *Chrysopa* | *oculata* | NorthAmerica | | -0.163 | 0.153 |
| Mean body size | Coleoptera | *Acanthoscelides* | *pallidipennis* | Asia | | -0.379 | 0.416 |
| Mean body size | Coleoptera | *Harmonia* | *axyridis* | Asia | | 0.01 | 0.295 |

| | | | | | | | |
|---|---|---|---|---|---|---|---|
| Mean body size | Coleoptera | *Leptinotarsa* | *decemlineata* | Europe | | -0.005 | 0.19 |
| Mean body size | Coleoptera | *Leptinotarsa* | *decemlineata* | NorthAmerica | | 1.481 | 0.065 |
| Mean body size | Diptera | *Aedes* | *albopictus* | NorthAmerica | | 0.265 | 0.038 |
| Mean body size | Diptera | *Drosophila* | *melanogaster* | Australia | | 0.553 | 0.048 |
| Mean body size | Diptera | *Scathophaga* | *stercoraria* | Europe | | 0.136 | 0.049 |
| Mean body size | Diptera | *Wyeomyia* | *smithii* | NorthAmerica | | -0.444 | 0.167 |
| Mean body size | Hemiptera | *Nezara* | *viridula* | Australia | | 3.004 | 1.539 |
| Mean body size | Lepidoptera | *Inachis* | *io* | Europe | | 0.415 | 0.946 |
| Mean body size | Lepidoptera | *Lasiommata* | *megera* | Europe | | -2.756 | 0.539 |
| Mean body size | Lepidoptera | *Ostrinia* | *furnacalis* | Asia | | -0.325 | 0.068 |
| Mean body size | Lepidoptera | *Pararge* | *aegeria* | Europe | | 0.18 | 0.054 |
| Mean body size | Lepidoptera | *Pieris* | *rapae* | Asia | | -0.66 | 0.432 |
| Mean body size | Lepidoptera | *Pieris* | *rapae* | Europe | | -0.18 | 0.107 |
| Mean body size | Lepidoptera | *Pieris* | *rapae* | NorthAmerica | | -0.184 | 0.145 |
| Mean body size | Lepidoptera | *Sericinus* | *montelus* | Asia | | -2.066 | 0.302 |
| Mean development rate | Coleoptera | *Acanthoscelides* | *pallidipennis* | Asia | total | 0.244 | 0.156 |
| Mean development rate | Coleoptera | *Bruchidius* | *dorsalis* | Asia | total | 0.1 | 0.382 |
| Mean development rate | Coleoptera | *Epicauta* | *gorhami* | Asia | larva | 2.087 | 0.787 |
| Mean development rate | Coleoptera | *Epicauta* | *gorhami* | Asia | pupa | 1.409 | 0.658 |
| Mean development rate | Coleoptera | *Harmonia* | *axyridis* | Asia | total | 0.134 | 0.074 |
| Mean development rate | Coleoptera | *Leptinotarsa* | *decemlineata* | Europe | total | 1.402 | 0.363 |
| Mean development rate | Coleoptera | *Leptinotarsa* | *decemlineata* | NorthAmerica | total | -0.119 | 0.04 |
| Mean development rate | Diptera | *Aedes* | *albopictus* | NorthAmerica | total | 0.228 | 0.055 |
| Mean development rate | Diptera | *Aedes* | *sierrensis* | NorthAmerica | larva | 0.244 | 0.219 |
| Mean development rate | Diptera | *Aedes* | *sierrensis* | NorthAmerica | pupa | 1.258 | 0.249 |
| Mean development rate | Diptera | *Drosophila* | *melanogaster* | Europe | total | -0.03 | 0.043 |
| Mean development rate | Diptera | *Scathophaga* | *stercoraria* | Europe | total | 0.186 | 0.069 |
| Mean development rate | Diptera | *Wyeomyia* | *smithii* | NorthAmerica | total | -0.797 | 0.231 |

| | | | | | | | |
|---|---|---|---|---|---|---|---|
| Mean development rate | Hemiptera | *Laodelphax* | *striatellus* | Asia | larva | -1.958 | 0.08 |
| Mean development rate | Hemiptera | *Nezara* | *viridula* | Australia | larva | -2.685 | 1.202 |
| Mean development rate | Hemiptera | *Rhopalosiphum* | *padi* | Asia | larva | 1.415 | 0.154 |
| Mean development rate | Lepidoptera | *Diatraea* | *grandiosella* | NorthAmerica | larva | -2.643 | 0.528 |
| Mean development rate | Lepidoptera | *Inachis* | *io* | Europe | total | -0.898 | 0.653 |
| Mean development rate | Lepidoptera | *Lasiommata* | *megera* | Europe | larva | -0.674 | 0.442 |
| Mean development rate | Lepidoptera | *Ostrinia* | *furnacalis* | Asia | total | 0.184 | 0.029 |
| Mean development rate | Lepidoptera | *Papilio* | *memnon* | Asia | egg | -0.488 | 0.44 |
| Mean development rate | Lepidoptera | *Papilio* | *memnon* | Asia | larva | -1.326 | 0.395 |
| Mean development rate | Lepidoptera | *Papilio* | *memnon* | Asia | pupa | -0.218 | 0.422 |
| Mean development rate | Lepidoptera | *Pararge* | *aegeria* | Europe | larva | 0.188 | 0.065 |
| Mean development rate | Lepidoptera | *Pararge* | *aegeria* | Europe | pupa | -0.44 | 0.051 |
| Mean development rate | Lepidoptera | *Pieris* | *napi* | Europe | egg | 1.215 | 0.199 |
| Mean development rate | Lepidoptera | *Pieris* | *napi* | Europe | larva | 0.231 | 0.239 |
| Mean development rate | Lepidoptera | *Pieris* | *napi* | Europe | pupa | 0.808 | 0.223 |
| Mean development rate | Lepidoptera | *Pieris* | *rapae* | Asia | total | -1.94 | 0.418 |
| Mean development rate | Lepidoptera | *Pieris* | *rapae* | NorthAmerica | total | 0.12 | 0.115 |
| Mean development rate | Lepidoptera | *Zygaena* | *trifolii* | Europe | total | -1.007 | 0.186 |
| Mean development rate | Neuroptera | *Chrysopa* | *oculata* | NorthAmerica | total | -0.04 | 0.256 |
| Body size plasticity | Diptera | *Aedes* | *albopictus* | NorthAmerica | | -0.02 | 0.063 |
| Body size plasticity | Diptera | *Scathophaga* | *stercoraria* | Europe | | 0.08 | 0.09 |
| Body size plasticity | Diptera | *Wyeomyia* | *smithii* | NorthAmerica | | 0.151 | 0.382 |
| Body size plasticity | Hemiptera | *Nezara* | *viridula* | Australia | | 4.423 | 8.15 |
| Body size plasticity | Lepidoptera | *Inachis* | *io* | Europe | | 0.02 | 1.729 |
| Body size plasticity | Lepidoptera | *Lasiommata* | *megera* | Europe | | 2.234 | 1.541 |
| Body size plasticity | Lepidoptera | *Ostrinia* | *furnacalis* | Asia | | -0.019 | 0.215 |
| Body size plasticity | Lepidoptera | *Pieris* | *rapae* | Asia | | -3.077 | 1.095 |
| Body size plasticity | Lepidoptera | *Pieris* | *rapae* | NorthAmerica | | 1.266 | 0.305 |

| | | | | | | | |
|---|---|---|---|---|---|---|---|
| Development rate plasticity | Coleoptera | *Epicauta* | *gorhami* | Asia | larval | 2.042 | 1.329 |
| Development rate plasticity | Coleoptera | *Epicauta* | *gorhami* | Asia | pupa | -0.52 | 1.758 |
| Development rate plasticity | Diptera | *Aedes* | *albopictus* | NorthAmerica | total | 0.407 | 0.06 |
| Development rate plasticity | Diptera | *Aedes* | *sierrensis* | NorthAmerica | larval | 0.12 | 0.239 |
| Development rate plasticity | Diptera | *Aedes* | *sierrensis* | NorthAmerica | pupa | -0.19 | 0.293 |
| Development rate plasticity | Diptera | *Drosophila* | *melanogaster* | Europe | total | -0.036 | 0.057 |
| Development rate plasticity | Diptera | *Scathophaga* | *stercoraria* | Europe | total | -0.279 | 0.121 |
| Development rate plasticity | Diptera | *Wyeomyia* | *smithii* | NorthAmerica | total | -0.991 | 0.327 |
| Development rate plasticity | Hemiptera | *Laodelphax* | *striatellus* | Asia | larval | 0.846 | 0.125 |
| Development rate plasticity | Hemiptera | *Nezara* | *viridula* | Australia | larval | 1.647 | 1.832 |
| Development rate plasticity | Lepidoptera | *Diatraea* | *grandiosella* | NorthAmerica | larval | -2.475 | 0.697 |
| Development rate plasticity | Lepidoptera | *Inachis* | *io* | Europe | total | 0.001 | 0.876 |
| Development rate plasticity | Lepidoptera | *Lasiommata* | *megera* | Europe | larval | 0.241 | 0.746 |
| Development rate plasticity | Lepidoptera | *Ostrinia* | *furnacalis* | Asia | total | 0 | 0.052 |
| Development rate plasticity | Lepidoptera | *Pararge* | *aegeria* | Europe | larval | 10.877 | 9.789 |
| Development rate plasticity | Lepidoptera | *Pararge* | *aegeria* | Europe | pupa | 8.229 | 4.297 |
| Development rate plasticity | Lepidoptera | *Pieris* | *napi* | Europe | egg | 1.775 | 0.335 |
| Development rate plasticity | Lepidoptera | *Pieris* | *napi* | Europe | larval | -0.189 | 0.214 |
| Development rate plasticity | Lepidoptera | *Pieris* | *napi* | Europe | pupa | 0.114 | 0.252 |
| Development rate plasticity | Lepidoptera | *Pieris* | *rapae* | Asia | total | -2.37 | 0.683 |
| Development rate plasticity | Lepidoptera | *Pieris* | *rapae* | NorthAmerica | total | 0.295 | 0.18 |
| Development rate plasticity | Neuroptera | *Chrysopa* | *oculata* | NorthAmerica | total | -0.375 | 0.41 |
